# Microbial nanowires with genetically modified peptide ligands to sustainably fabricate electronic sensing devices

**DOI:** 10.1101/2022.10.17.512539

**Authors:** Yassir Lekbach, Toshiyuki Ueki, Xiaomeng Liu, Trevor Woodard, Jun Yao, Derek R. Lovley

## Abstract

Nanowires have substantial potential as the sensor component in electronic sensing devices. However, surface functionalization of traditional nanowire and nanotube materials with short peptides that increase sensor selectivity and sensitivity requires complex chemistries with toxic reagents. In contrast, microorganisms can assemble pilin monomers into protein nanowires with intrinsic conductivity from renewable feedstocks, yielding an electronic material that is robust and stable in applications, but also biodegradable. Here we report that the sensitivity and selectivity of protein nanowire-based sensors can be modified with a simple plug and play genetic approach in which a short peptide sequence, designed to bind the analyte of interest, is incorporated into the pilin protein that is microbially assembled into nanowires. We employed a scalable *Escherichia coli* chassis to fabricate protein nanowires that displayed either a peptide previously demonstrated to effectively bind ammonia, or a peptide known to bind acetic acid. Sensors comprised of thin films of the nanowires amended with the ammonia-specific peptide had a ca. 100-fold greater response to ammonia than sensors made with unmodified protein nanowires. Protein nanowires with the peptide that binds acetic acid yielded a 4-fold higher response than nanowires without the peptide. The results demonstrate that protein nanowires with enhanced sensor response for analytes of interest can be fabricated with a flexible genetic strategy that sustainably eliminates the energy, environmental, and health concerns associated with other common nanomaterials.

## 1. Introduction

Nanowires are desirable electronic materials because they facilitate miniaturization and convey flexibility to electronics. They are particularly important for fabricating electronic sensors with improved sensing performance (Patolsky and Lieber 2005). Adding functional groups to the nanowire surface can lead to specific binding of analytes of interest for more selective detection. However, the traditional chemistries for attaching functional groups are complex. Furthermore, common non-biological synthetic materials such as silicon nanowires and carbon nanotubes pose serious sustainability challenges due to requirements for toxic chemicals and/or high energy inputs for synthesis. High temperatures are required to generate silicon nanowires and carbon nanotubes and fabrication of silicon nanowires also requires the vaporization of highly toxic components (Hu et al. 1999; Prasek et al. 2011). The need for a clean-room environment for material production increases costs and technical complexity, limiting the feasibility of mass production. These non-biological nanomaterials are not biodegradable and carbon nanotubes are toxic and carcinogenic (Hansen and Lennquist 2020).

In contrast, microorganisms can sustainably produce non-toxic electrically conductive protein nanowires from renewable organic feedstocks (Lovley 2017; Lovley and Yao 2021). Most notable are the 3 nm diameter conductive protein nanowires assembled from the native pilin protein of *Geobacter sulfurreducens* (Lovley 2022a, b). These pilin-based protein nanowires have served as the electronic components in a diversity of applications including: devices that generate electricity from atmospheric humidity (Liu et al. 2020b); neuromorphic memory devices (Fu et al. 2021; Fu et al. 2020b); and sensors (Liu et al. 2020a; Smith et al. 2020). A key feature of pilin-based nanowires is that their function can readily be modified with simple changes to the pilin gene sequence. Pilin-based nanowire conductivity was tuned over a million-fold (40 µS/cm to 277 S/cm at pH 7) simply by modifying the pilin gene sequence to adjust the abundance of aromatic amino acids in the pilin protein (Adhikari et al. 2016; Liu et al. 2021; Tan et al. 2017; Tan et al. 2016). In addition to their ‘green’ synthesis, pilin-based nanowires are robust with long-term stability in electronics applications (Liu et al. 2020a; Liu et al. 2020b; Smith et al. 2020), but are also biodegradable, avoiding the accumulation of electronic waste (Lovley 2017; Lovley and Yao 2021).

Sensors that can detect volatile compounds have broad biomedical and environmental applications (Ge et al. 2020; Rasheed et al. 2020). Vapor sensor designs often rely on pattern recognition algorithms to interpret the binding of analytes to sensor arrays, but a more direct sensing approach is to design sensor elements that specifically bind analytes of interest (Barbosa et al. 2018; McAlpine et al. 2008; Wasilewski et al. 2022). Peptides can be designed to function as ligands for specific chemical and biological targets (Pardoux et al. 2020; Sfragano et al. 2021; Wu et al. 2001). For example, guidance from the binding domains of human olfactory receptor proteins, coupled with molecular simulations and experimental verification, has identified peptides that specifically bind gases of interest (Wu et al. 2001). Silicon nanowires (McAlpine et al. 2008) and carbon nanotubes (Li et al. 2020; Palomar et al. 2020) can be functionalized with peptides to improve selectivity of nanowire-based sensors, but in addition to the limitations noted above in producing the nanowire material, the peptide sensor components have to be synthesized and purified in an expensive complex process requiring toxic reagents.

In contrast, decorating pilin-based protein nanowires with desired peptide sequences is sustainably achieved with simple and versatile modifications to the pilin gene sequence (Ueki et al. 2019). Pilin gene sequences customized to encode 6-9 extra amino acids at the carboxyl end of the pilin yielded nanowires in which the added amino acid sequences were displayed along the outer surface of the nanowire without interfering with nanowire conductivity. This approach offers a strategy for displaying peptide ligands on the outer surface of nanowires for potential sensing applications that is much more programable and sustainable than the methods for functionalizing non-biological nanowire materials.

Therefore, we investigated whether decorating pilin-based protein nanowires with peptides designed to bind analytes of interest could increase the sensing response obtained in pilin-based electronic gas sensors. We focused on ammonia and acetic acid analytes, which were also the focus of similar studies with silicon nanowires (McAlpine et al. 2008) because these volatiles in breath are indicators of kidney disease (ammonia) (Ricci and Gregory 2021) and asthma (acetic acid) (Pineau et al. 2021). We expressed the customized protein nanowires in an *Escherichia coli* chassis engineered to assemble nanowires from the *G. sulfurreducens* pilin gene (Ueki et al. 2020). This approach provides a simple method for mass production of pilin-based nanowires while avoiding the possibility that the nanowire preparations are contaminated with other *G. sulfurreducens* outer surface proteins (Ueki et al. 2020). The results demonstrate that pilin-based nanowires can be designed to specifically enhance sensor response to analytes of interest.

## 2. Material and methods

### 2.1 Construction of E. coli strains for nanowire expression

*E. coli* strains for the production of nanowires for sensing ammonia or acetic acid were constructed as described previously (Ueki et al. 2020) with modifications as follows. The *G. sulfurreducens* pilin gene was extended to encode peptides that were previously found (McAlpine et al. 2008; Wu et al. 2001) to specially bind either ammonia (DLESFL) or acetic acid (RVNEWVI) at the carboxyl end of the pilin protein. DNA fragments for the nanowire monomers for ammonia or acetic acid were amplified with the PCR with primer pairs, Gs*pilA*-F (TCTCATATGGACAAGCAACGCGGTTTCACCCTTATCGAGCTGC)/Gs*pilA*-Am-R (TCTGAGCTCTTACAGAAAGCTCTCCAGATCACTTTCGGGCGGATAGGTTTG) or Gs*pilA*-F (TCTCATATGGACAAGCAACGCGGTTTCACCCTTATCGAGCTGC)/Gs*pilA*-Ac-R (TCTGAGCTCTTAGATAACCCACTCATTAACGCGACTTTCGGGCGGATAGGTTTG), respectively. The amplified DNA fragments were digested with NdeI and SacI and then cloned into the nanowire expression vector T4PAS/p24Ptac (Ueki et al. 2020). The resultant plasmids, designated Gs*pilA*-AMM/T4PAS/p24Ptac (ammonia) or Gs*pilA*-ACE/T4PAS/p24Ptac (acetic acid), were transformed into *E. coli* Δ*fimA*Δ*fliC*, a strain in which genes for FimA, the primary monomer for type I pili, and FliC, the structural flagellin of flagella, were deleted. Strain Δ*fimA*Δ*fliC* (kanamycin-sensitive) was constructed by deleting the *fliC* gene from strain Δ*fimA* (Ueki et al., 2020) as described previously (Baba et al. 2006; Datsenko and Wanner 2000). The amino acid sequences of the unmodified pilin, the pilin with the ammonia-binding peptide, and the pilin with the acetic acid-binding peptide were:

Unmodified pilin:

FTLIELLIVVAIIGILAAIAIPQFSAYRVKAYNSAASSDLRNLKTALESAFADDQTYPPES

Pilin modified with ammonia-binding peptide:

FTLIELLIVVAIIGILAAIAIPQFSAYRVKAYNSAASSDLRNLKTALESAFADDQTYPPESDLESFL

Pilin modified with acetic acid-binding peptide:

FTLIELLIVVAIIGILAAIAIPQFSAYRVKAYNSAASSDLRNLKTALESAFADDQTYPPESRVNEWVI

### 2.2 Protein nanowire fabrication

*E. coli* strains were grown aerobically at 30 °C in agar-solidified LB medium supplemented with kanamycin (50 µg/ml) (Ueki et al. 2020). After 24 h incubation, the cells were gently scraped off the agar and then spread plated onto agar-solidified M9 medium held in sterile stainless steel trays (37 cm × 27 cm × 6 cm). M9 medium consists of Na_2_HPO_4_-7H_2_O, 12.8 g/l; KH_2_PO_4_, 3 g/l; NaCl, 0.5 g/l; NH_4_Cl, 1 g/l; MgSO_4_, 2 mM; CaCl_2_, 0.1 mM; glycerol, 0.5%,; IPTG, 0.5 mM; kanamycin, 50 µg/ml and agar 15 g/l. After 48 h incubation at 30 °C, bacterial cells were scraped from the agar surface and suspended in M9 medium. The suspension was centrifuged to harvest cells, and the resultant pellets were suspended in ethanolamine HCl buffer (150 mM, pH 10.5). Protein nanowires were purified with an ammonium sulfate precipitation method, as previously described (Liu et al. 2020b). Briefly, protein nanowires were sheared from the bacterial suspension in a blender at low speed. The resultant solution was centrifuged to remove cell debris. The protein nanowires in the supernatant were precipitated with ammonium sulfate (20%), followed by centrifugation, and then resuspended in ethanolamine HCl (150 mM, pH 10.5). Impurities were removed with a 1% ammonium sulfate precipitation and subsequent centrifugation. Protein nanowires were precipitated in 18% ammonium sulfate and collected via centrifugation. Pellets were suspended in ethanolamine HCl (150 mM, pH 10.5) and then dialyzed against deionized water to remove salts. The purified nanowires were suspended in 2 ml of sterile water and stored at 4°C until use. Protein concentration was determined using the BCA protein assay kit (Thermo Pierce, USA) according to the manufacturer’s instructions.

### 2.3 Sensor construction

The gas sensing devices were prepared as previously described (Smith et al. 2020). Briefly, a pair of interdigitated electrodes was fabricated on a Si/SiO_2_ wafer with standard lithography, metal deposition (Cr/Au, 5/50 nm), and lift-off processes. The width of each electrode was 400 µm and the electrode separation was 100 µm. Ten µl of a suspension of purified protein nanowires solution (70 µg/ml) were drop-casted onto the surface of the pair of interdigitated electrodes and left to dry at room temperature.

The sensor was connected to a semiconductor characterization system (Keithley 4200-SCS) and placed inside a custom-built airtight test chamber (Fig. 1). A voltage of 1 V was applied across the electrodes. An air pump provided a steady stream of air that entered the test chamber through a tubing connection. The relative humidity of the air was constant (21 ± 1 %) throughout the testing process. Vapor samples to be evaluated were injected into the air stream through a septum with a syringe and needle.

**Fig. 1.**
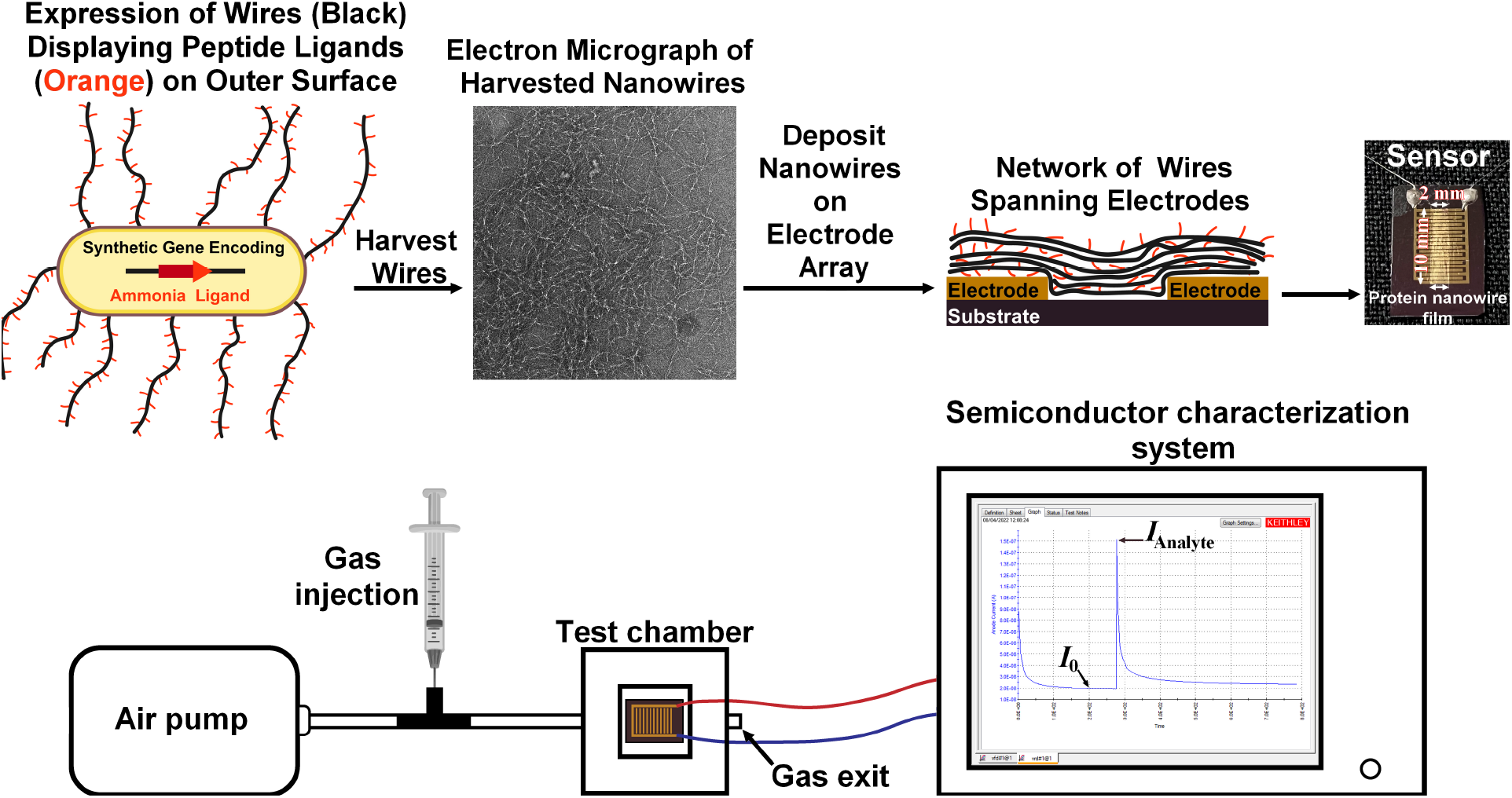
Schematic of sensor fabrication and evaluation.

The sensor responses were calculated using the following formula (Chou et al. 2018; Jha et al. 2018):

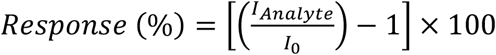

where *I*_0_ was the background current measured when just air was passing through the system and *I*_*Analyte*_ was the maximum current when the gas sample passed through the test chamber.

## 3. Results and Discussion

Preparations of outer surface filaments harvested from *G. sulfurreducens*, which are dominated by pilin-based nanowires (Fu et al. 2020b; Liu et al. 2021), effectively functioned as the sensor element for specifically detecting ammonia, but not other gases typically present in human breath, such as carbon dioxide, ethanol, or acetone (Smith et al. 2020). In an effort to increase the response to ammonia, the *G. sulfurreducens* pilin gene was modified to encode the peptide DLESFL, which has a high affinity for ammonia gas (Wu et al. 2001), at the carboxyl terminus of the pilin. Prior studies have indicated that the added peptide can be expected to be displayed on the outer surface of the microbially assembled nanowires (Ueki et al. 2019), thus providing ligands for ammonia along the length of the nanowires. The pilin gene was expressed in *E. coli* to avoid the possibility of contamination of the protein nanowire preparation by other nanofilaments expressed by *G. sulfurreducens* (Ueki et al. 2020).

As expected from previous studies with pili produced with *G. sulfurreducens* (Smith et al. 2020), the nanowires that *E. coli* assembled from the unmodified *G. sulfurreducens* pilin responded to ammonia with increasing current output as ammonia concentrations increased (Fig. 2a-c). The current output from devices with an equivalent quantity of nanowires customized with the ammonia-binding peptide was ca. 100-fold higher than the output from the nanowires assembled from the unmodified pilin (Fig. 2d-f, Fig. 3a,b). The response to ammonia was rapid and the electrical signal quickly returned to baseline as the air flow flushed the ammonia from the sensing chamber. These results demonstrated that modifying the nanowires with the ammonia ligand substantially enhanced the response to ammonia and suggested that ammonia binding to the ligand was readily reversible as the ammonia was rapidly re-released into the overlying air stream. Thus, the sensor is capable of detecting dynamic changes in ammonia concentrations in real time.

**Fig. 2.**
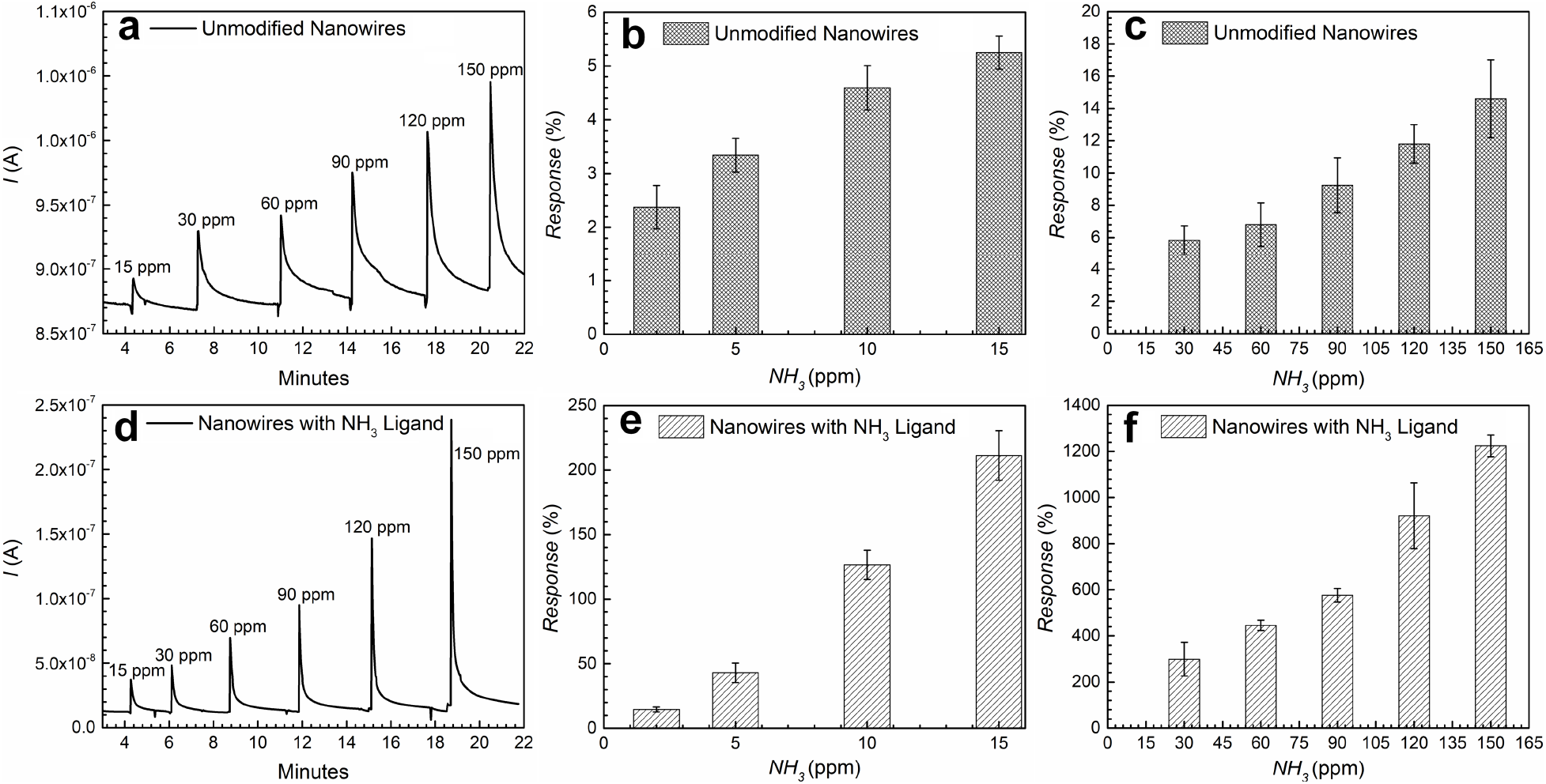
Current outputs in response to injections of different ammonia concentrations in (a-c) sensor devices with nanowires assembled from unmodified pilin or (d-f) sensor devices with nanowires assembled from pilin with ammonia-specific peptide. Data in panels a and d are representative current outputs from triplicate sensing devices. Bars and error bars in panels b,c,e, and f designate the means and standard deviations from triplicate sensor devices.

**Fig. 3.**
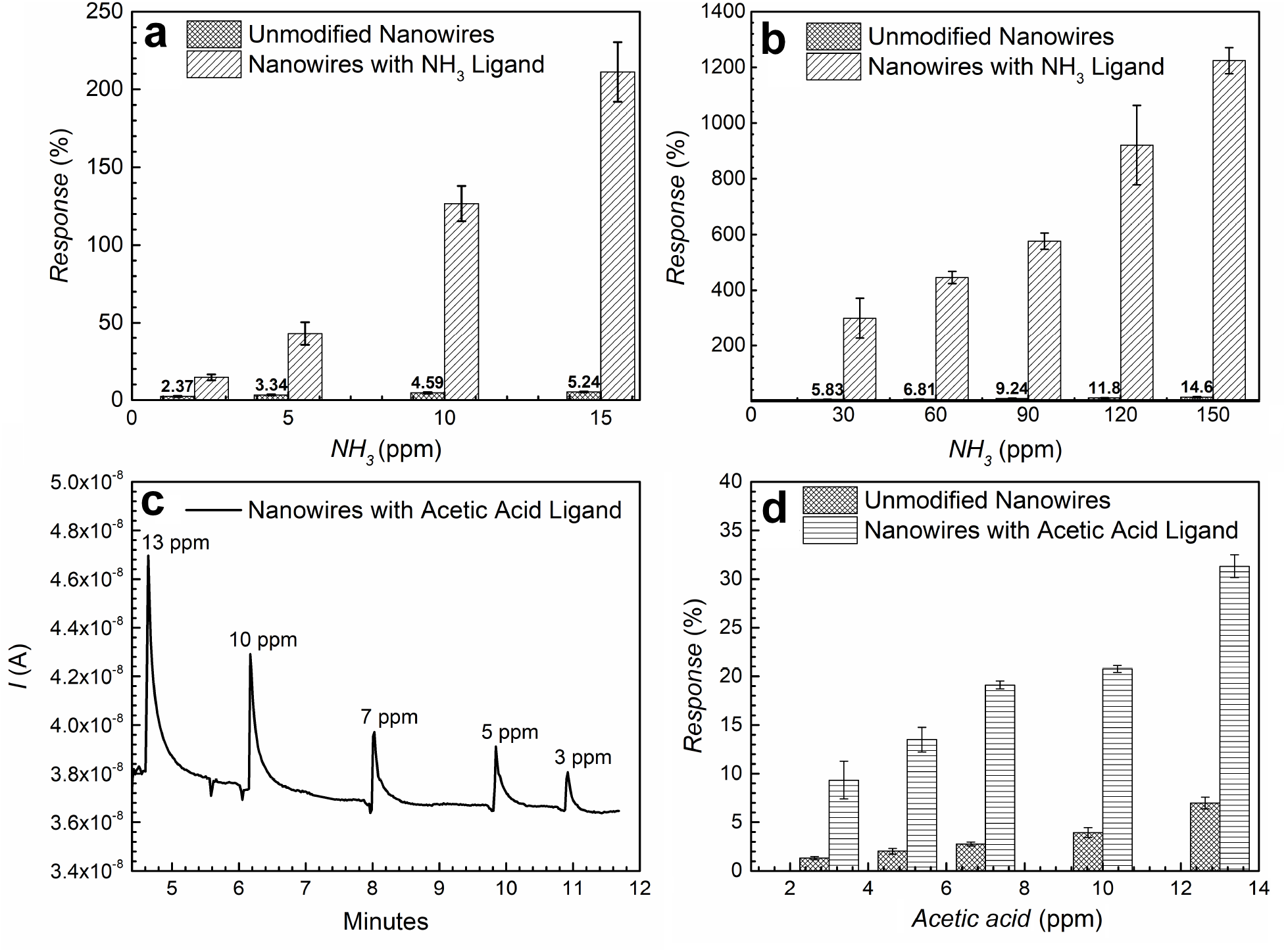
Relative current response of unmodified nanowires and nanowires with analyte-specific ligands. (a, b) Greater response of nanowires modified with an ammonia-specific peptide to ammonia. (c) Representative current output in response to injections of acetic acid in sensor with nanowires assembled from pilin with the acetic acid-specific peptide. (d) Greater response of nanowires modified with an acetic acid-specific peptide to acetic acid. Bars and error bars designate the means and standard deviations from triplicate sensor devices.

The peptide RVNEWVI has a high affinity for acetic acid (Wu et al. 2001). A pilin gene which encoded the RVNEWVI amino acid sequence at the carboxyl terminus yielded nanowires with a rapid response to acetic acid (Fig. 3c) that was ca. 4-fold higher than sensors fabricated with the unmodified nanowires (Fig. 3d). Although the relative increase in current output achieved with the acetic acid ligand modification was smaller than that with the ammonia-specific ligand, the results do further demonstrate that nanowires can be customized to improve sensor response.

The ligand additions selectively increased response to the intended analyte. The current response to 13 ppm acetic acid for sensor devices fabricated with the nanowires modified with the ammonia-specific ligand (6.51±0.76%; mean ± standard deviation, n=3) was similar to the response with unmodified nanowires (6.98±0.61%), confirming the specificity of these modified nanowires for sensing ammonia. This result is consistent with the previous finding that the DLESFL peptide has a much higher affinity for ammonia than acetic acid with a selectivity ratio of 75:1 (McAlpine et al. 2008).

In previous studies the selectivity of the RVNEWVI peptide for acetic acid versus ammonia was only 3.75:1 (McAlpine et al. 2008). In accordance with these findings, the nanowires modified with RVNEWVI to enhance acetic acid binding had a higher response to ammonia at 150 ppm (44.2 ± 5.05%) than the unmodified nanowires (14.6 ± 2.41%). However, the increased response of the nanowires modified with RVNEWVI was much less than the response to 150 ppm ammonia (1224 ± 47.2%) of the nanowires modified with the DLESFL peptide designed for binding ammonia.

## 4. Conclusions

The results demonstrate that pilin-based protein nanowires for sensor applications can be fabricated with an *E. coli* chassis and that the sensing response of the pilin-based nanowires can be genetically tuned for higher sensitivity by incorporating specific amino acid sequences at the carboxyl end of the pilin monomer. This simple, low energy, ‘green’ synthesis of peptide-functionalized nanowire sensing components is in marked contrast to the fabrication of non-biological nanowire materials, which require complex fabrication procedures that involve high energy inputs and toxic chemicals and/or yield toxic products. Previous studies have demonstrated that it is possible to express individual protein nanowires with multiple different peptide ligands and to control the stoichiometry of ligand display along the length of the protein nanowires with precise control over genetic expression circuits (Ueki et al. 2019). This further expands the sensor design possibilities beyond what is readily possible with non-biological nanowire materials.

Peptides have been designed to specifically bind other volatiles, such as aldehydes (Wasilewski et al. 2018), trimethylamine (Lee et al. 2015), isopropyl alcohol, isoprene, toluene (Sankaran et al. 2011), o-xylene, (Wu et al. 2001) butyric acid, dimethyl amine, benzene, and chlorobenzene (Lu et al. 2009). Thus, microbially produced nanowires might be designed for effective sensing of a diversity of gases of biomedical, environmental, or practical importance. It may also be possible to tailor protein nanowires for sensing non-volatiles such as proteins (Vanova et al. 2021), viruses (Fu et al. 2020a), pathogenic bacteria (Bruce and Clapper 2020; Pardoux et al. 2019), and metallic ions (Liu et al. 2015; Ramezanpour et al. 2021).

These possibilities combined with potential to power protein nanowire sensors with protein nanowire-based devices that harvest electricity from atmospheric humidity (Fu et al. 2021; Liu et al. 2020b), or biofilm devices that generate electricity from sweat evaporation (Liu et al. 2022), coupled with protein nanowire-based devices to interpret the sensor outputs (Fu et al. 2021; Fu et al. 2020b), demonstrate the many opportunities for developing sustainable, self-powered monitoring devices for biomedical and environmental applications.

## Acknowledgement

J.Y. and D.R.L. acknowledge support from the National Science Foundation (NSF) DMR2027102.

## Notes

### Competing Interest Statement

The authors have declared no competing interest.

